# Profiling the Landscape of Drug Resistance Mutations in Neosubstrates to Molecular Glue Degraders

**DOI:** 10.1101/2021.12.23.474016

**Authors:** Pallavi M. Gosavi, Kevin C. Ngan, Megan Yeo, Cindy Su, Jiaming Li, Nicholas Z. Lue, Samuel M. Hoenig, Brian B. Liau

## Abstract

Targeted protein degradation (TPD) holds immense promise for drug discovery but mechanisms of acquired resistance to degraders remain to be fully identified. Here we used CRISPR-suppressor scanning to identify mechanistic classes of drug resistance mutations to molecular glue degraders in GSPT1 and RBM39, neosubstrates targeted by E3 ligase substrate receptors cereblon and DCAF15, respectively. While many mutations directly alter the ternary complex heterodimerization surface, distal resistance sites were also identified. Several distal mutations in RBM39 led to modest decreases in degradation yet can enable cell survival, underscoring how small differences in degradation can lead to resistance. Integrative analysis of resistance sites across GSPT1 and RBM39 revealed varying levels of sequence conservation and mutational constraint that control the emergence of different resistance mechanisms, highlighting that many regions co-opted by TPD are inessential. Altogether, our study identifies common resistance mechanisms for molecular glue degraders and outlines a general approach to survey neosubstrate requirements necessary for effective degradation.

## Introduction

In recent years, the discovery of molecular glue degraders has converged with the development of proteolysis targeting chimeras (PROTACs), revealing the remarkable ability of small molecules to co-opt the ubiquitin-proteasome system (UPS) and degrade protein targets.^1–4^ Molecular glue degraders chemically remodel E3 ligase substrate receptors, creating a small molecule-protein composite surface capable of *de novo* complexation with complementary yet otherwise unrelated protein substrates. These neosubstrates are then subsequently polyubiquitinated and proteolytically degraded via the UPS.^5^

Immunomodulatory drugs (IMiDs), including thalidomide and its analogs lenalidomide and pomalidomide, bind to cereblon (CRBN), a substrate receptor for the CUL4-RING (CRL4) E3 ubiquitin ligase, and induce its complexation with various neosubstrates that are subsequently degraded.^6–9^ Mechanistic studies of IMiDs revealed that the selectivity of the CRBN-IMiD recognition surface and their targeted neosubstrates could be broadly modulated through even subtle chemical changes to the IMiD structure.^10,11^ As a leading example, CC-885 (**Fig 1a**), an analog of lenalidomide, was shown to gain the ability to induce degradation of GSPT1 (also known as eRF3A), a translation termination factor essential for acute myeloid leukemia (AML) cell proliferation.^10^ Structural studies on the CC-885-CRBN-GSPT1^11,12^ ternary complex were critical in determining a β-hairpin structural degron, a unifying motif across the diverse array of IMiD-targeted neosubstrates necessary for CRL4^CRBN^-mediated degradation.^10,12,13^ New IMiD derivatives tailored to degrade novel neosubstrates, including GSPT1 and IKZF2, have entered clinical trials for oncology applications.

**Figure 1.**
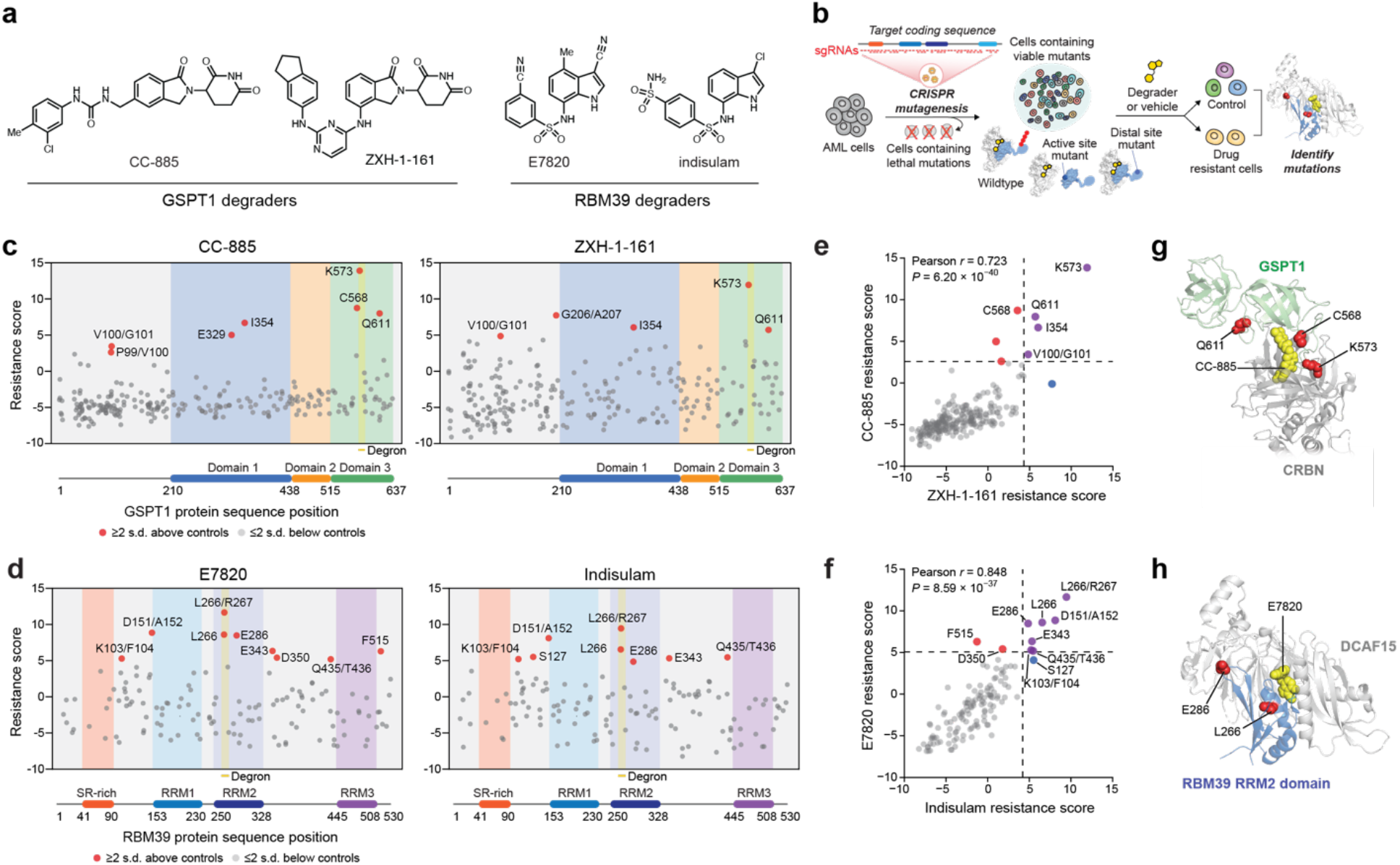
CRISPR-suppressor scanning identifies regions of GSPT1 and RBM39 that mediate targeted protein degradation by molecular glue degraders. a) Chemical structures of CC-885 and E7820. b) Schematic showing the CRISPR-suppressor scanning workflow applied to molecular glue degraders. c) Scatter plot showing resistance scores (*y* axis) in MOLM-13 under CC-885 (left) or ZXH-1-161 (right) treatment at 4 weeks. Resistance scores were calculated as the log_2_(fold-change sgRNA enrichment under drug treatment) normalized to the mean of the negative control sgRNAs (*n* = 22). The *GSPT1*-targeting sgRNAs (*n* = 239) are arrayed by amino acid position in the *GSPT1* CDS on the *x* axis corresponding to the position of the predicted cut site. When the sgRNA cut site falls between two amino acids, both amino acids are denoted. Data points represent mean values across three replicate treatments. Protein domains and the structural degron site are demarcated by the colored panels. d) Scatter plots showing resistance scores (*y* axis) in MOLM-13 under E7820 (left) or indisulam (right) treatment at 4 weeks. Resistance scores were calculated as the log_2_(fold-change sgRNA enrichment under drug treatment) normalized to the mean of the negative control sgRNAs (*n* = 77). The *RBM39*-targeting sgRNAs (*n* = 129) are arrayed by amino acid position in the *RBM39* CDS on the *x* axis corresponding to the position of the predicted cut site. Data points represent the mean values across three replicate treatments. Protein domains and the structural degron site are demarcated by the colored panels. e) Scatter plot showing *GSPT1*-targeting sgRNA resistance scores under CC-885 (*y* axis) or ZHX-1-161 (*x* axis) treatment at 4 weeks. Dotted lines represent 2 s.d. above the mean of the negative control sgRNAs. Pearson’s *r* and two-sided *P* value are shown. f) Scatter plot showing *RBM39*-targeting sgRNA resistance scores under E7820 (*y* axis) or indisulam (*x* axis) treatment at 4 weeks. Dotted lines represent 2 s.d. above the mean of the negative control sgRNAs. Pearson’s *r* and two-sided *P* value are shown. g) Structural view of the CC-885-CRBN-GSPT1 ternary complex showing the location of top-enriched sgRNAs (red) (Protein Data Bank (PDB): 5HXB). h) Structural view of the E7820-DCAF15-RBM39(RRM2) ternary complex showing the location of top-enriched sgRNAs (red) (PDB: 6UE5).

Aside from IMiDs, the anti-cancer sulfonamides, including E7820 and indisulam (**Fig 1a, Fig S1a**), were discovered to also operate as molecular glue degraders, highlighting the structural diversity of small molecules, neosubstrates, and E3 ligases that can be involved in TPD.^14–19^ These sulfonamides induce ternary complex formation between the CRL4 E3 substrate receptor DCAF15 and the splicing factors RBM39 (also known as CAPERα) and RBM23, which share a common α-helical structural degron.^14–19^ Notably, CRBN-IMiD and DCAF15-sulfonamide complexes engage their respective targets through completely distinct structural degrons and kinetic pathways,^17–19^ highlighting their unique modes of action despite thematic similarities. Taken together, the ability to co-opt the UPS and diverse E3 substrate receptors to induce degradation of a wide repertoire of unrelated protein targets – spanning transcription factors, kinases, translation regulators, and RNA-binding proteins – underscores TPD as a transformative approach for developing therapeutics against targets previously considered undruggable.^1,2,4^

As an emerging therapeutic modality, molecular glue degraders may encounter mechanisms of acquired resistance that differ substantially from canonical occupancy-driven inhibitors, potentially exploiting the unique molecular requirements necessary to catalyze proteolytic degradation.^2,4^ For instance, loss-of-function (LOF) and missense mutations in the IMiD-binding domain of CRBN confer resistance to IMiDs, which have been observed in multiple myeloma patients refractory to lenalidomide and pomalidomide.^4,20–22^ Additionally, multiple studies have shown that loss of other UPS components or chaperones can interfere with TPD.^23–29^ By contrast, systematic characterization of resistance mutations arising in the targeted neosubstrate has been more limited.^13,14^ Profiling the landscape of neosubstrate resistance mutations could delineate thematic classes of resistance mechanisms available to cancer cells and identify the structural and functional constraints that modulate their accessibility. More broadly, these mutational landscapes could illuminate molecular requirements and structural features – both within and beyond the structural degron – necessary for effective TPD.^13^ Motivated by these considerations, here we conducted CRISPR-suppressor scanning to systematically identify mutations across GSPT1 and RBM39 that confer resistance to molecular glue degraders, with the aim of investigating potentially unifying principles across distinct E3 substrate receptors and neosubstrates.

## Results

### CRISPR-suppressor scanning of *GSPT1* and *RBM39*

To identify candidate drug resistance mechanisms to molecular glue degraders, we performed CRISPR-suppressor scanning across two different TPD targets, GSPT1 and RBM39, which are recognized by distinct CRL4 substrate receptors. GSPT1 and RBM39 are both essential for the proliferation of AML cells and are clinical targets of interest for the treatment of AML.^30,31^ Consequently, we conducted CRISPR-mutagenesis of both *GSPT1* and *RBM39* in MOLM-13 cells, an *MLL*-rearranged AML cell line, to allow more facile comparisons across the two systems. In CRISPR-suppressor scanning,^32^ pools of single guide RNAs (sgRNAs) spanning either the *GSPT1* or *RBM39* coding sequence and control sgRNAs were transduced along with *Streptococcus pyogenes* Cas9 (SpCas9) into MOLM-13 cells (**Fig 1b**). DNA double-strand breaks introduced by Cas9 can lead to the formation of diverse insertion/deletion (indel) mutations at positions proximal to the sgRNA cut site. Cells containing lethal LOF mutations in either *GSPT1* or *RBM39* drop out, leaving pools of cells containing viable in-frame variants that are then split and treated with either vehicle or the appropriate molecular glue degraders to select for candidate drug resistance-conferring mutations (**Fig 1a, Fig S1a**).

For *GSPT1* CRISPR-suppressor scanning, the GSPT1 degraders CC-885 and ZXH-1-161^33^ were dosed in gradual escalation due to their acute cytotoxicity, starting at the approximate GI_50_ values and then gradually escalating to above the GI_90_ dose over 4 weeks (**Fig S1a**). In the case of *RBM39* CRISPR-suppressor scanning, E7820 (1 µM) and indisulam (1 µM) were dosed at the approximate GI_90_ concentrations for 4 weeks (**Fig S1b**). After vehicle or degrader treatment, genomic DNA was isolated from surviving cells and sequenced to deconvolute sgRNA identities enriched in each condition, allowing us to calculate “resistance scores” for each sgRNA that correspond to their enrichment in degrader-treated cells and hence their propensity to generate drug resistance-conferring mutations (**Fig 1c,d**). Enriched sgRNAs were asymmetrically distributed across the *GSPT1* and *RBM39* coding sequences in the degrader-treatment conditions, consistent with the expansion of drug-selected populations. For both neosubstrate targets, sgRNAs enriched in either set of drug treatments (CC-885 and ZXH-1-161 or E7820 and indisulam) were strongly correlated (**Fig 1e,f**), reflecting the structural similarity between the degraders employed and the overall assay robustness.

The top enriched sgRNAs across both screens targeted the structural degron of each respective TPD substrate (**Fig 1c,d**). For the GSPT1 screen, sgRNAs highly enriched in degrader treatment (i.e., sgC568, sgK573, sgQ611) clustered near the key β-hairpin structural degron and the CRBN-GSPT1 interface (**Fig 1g**).^10^ Likewise, top enriched sgRNAs in the RBM39 screen, sgL266 and sgL266/R267, target the α-helical structural degron in the RRM2 domain helix 1 that mediates the ternary DCAF15-RBM39-sulfonamide interaction (**Fig 1h**).^17–19^ These highly enriched sgRNAs presumably lead to mutations that disrupt ternary complex formation. However, several sgRNAs enriched in *RBM39*, and to a lesser extent, *GSPT1*, targeted positions distal to the structural degron that have not been previously implicated in degradation (i.e., RBM39 sgD151/A152, sgE286, sgE343). Altogether, these data demonstrate that CRISPR-suppressor scanning can identify key binding sites in neosubstrate targets previously established to be critical for ternary complex formation and, by extension, TPD.

### Identification of resistance mutations in the primary structural degrons

To investigate resistance mechanisms at a molecular level, we performed targeted amplicon deep sequencing directly from the pooled CRISPR-suppressor scan, focusing on the sgRNA cut sites corresponding to the highest enriched sgRNAs. This included sgRNAs targeting: (1) the region proximal to the β-hairpin structural degron of GSPT1 (**Fig 1g**, sgC568, sgK573, sgQ611), (2) the RRM2 helix 1 structural degron of RBM39 (**Fig 1h**, sgL266, sgL266/R267), and (3) the region N-terminal to the RRM1 domain of RBM39 (sgD151/A152). We genotyped *GSPT1* and *RBM39* variants and quantified their mutational allele frequency, which exhibited considerable variation in enrichment and sequence diversity (**Fig 2a, Fig 3a**).

**Figure 2.**
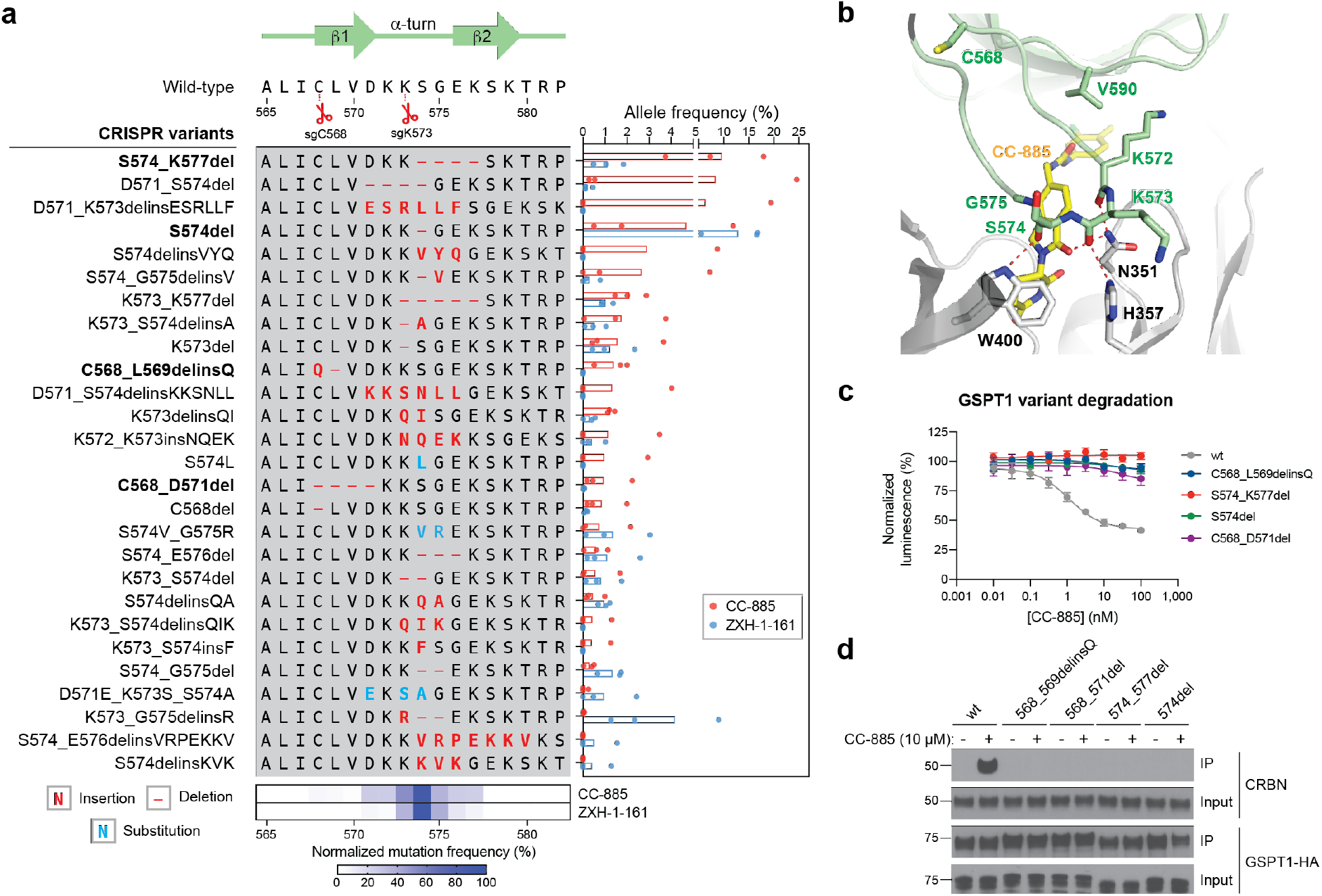
CC-885 resistance mutations alter the GSPT1 β-hairpin structural degron and impair GSPT1 degradation. a) Left: Schematic shows the coding variants of the most abundant in-frame mutations enriched in the β-hairpin structural degron of GSPT1 (>1% frequency in any condition). Right: Bar plot showing frequency (%, *x* axis) of each variant. Bars represent the mean across three replicate treatments and dots show the individual replicate values. Bottom: Heat map showing normalized mutational frequency (*y* axis, %) by sequence position (*x* axis). Mutational frequency was normalized as a percentage of the total frequency of the displayed variants. b) Structural view of the CC-885-CRBN-GSPT1 ternary complex, with key residues in CRBN (gray) and GSPT1 (green) highlighted. Carbon atoms of CC-885 are depicted in yellow. (PDB: 5HXB) c) Dose-response curves for wt and mutant HiBiT-GSPT1-HA cellular protein levels, as indicated by vehicle-normalized luminescence (*y* axis, %), in HEK293T cells treated with CC-885 for 6 h. Data represent mean ± s.e.m. across three technical replicates. One of two independent experiments is shown. d) Immunoblots showing co-IP of GSPT1-HA wt and mutant variants with CRBN after vehicle or CC-885 treatment (10 µM, 2 h) in transiently transfected HEK293T cells. All cells were first pre-treated with MLN-4924 (1 µM, 3 h) prior to vehicle or CC-885 treatment. Co-IP was performed using anti-HA antibody. One of two independent replicates is shown.

**Figure 3.**
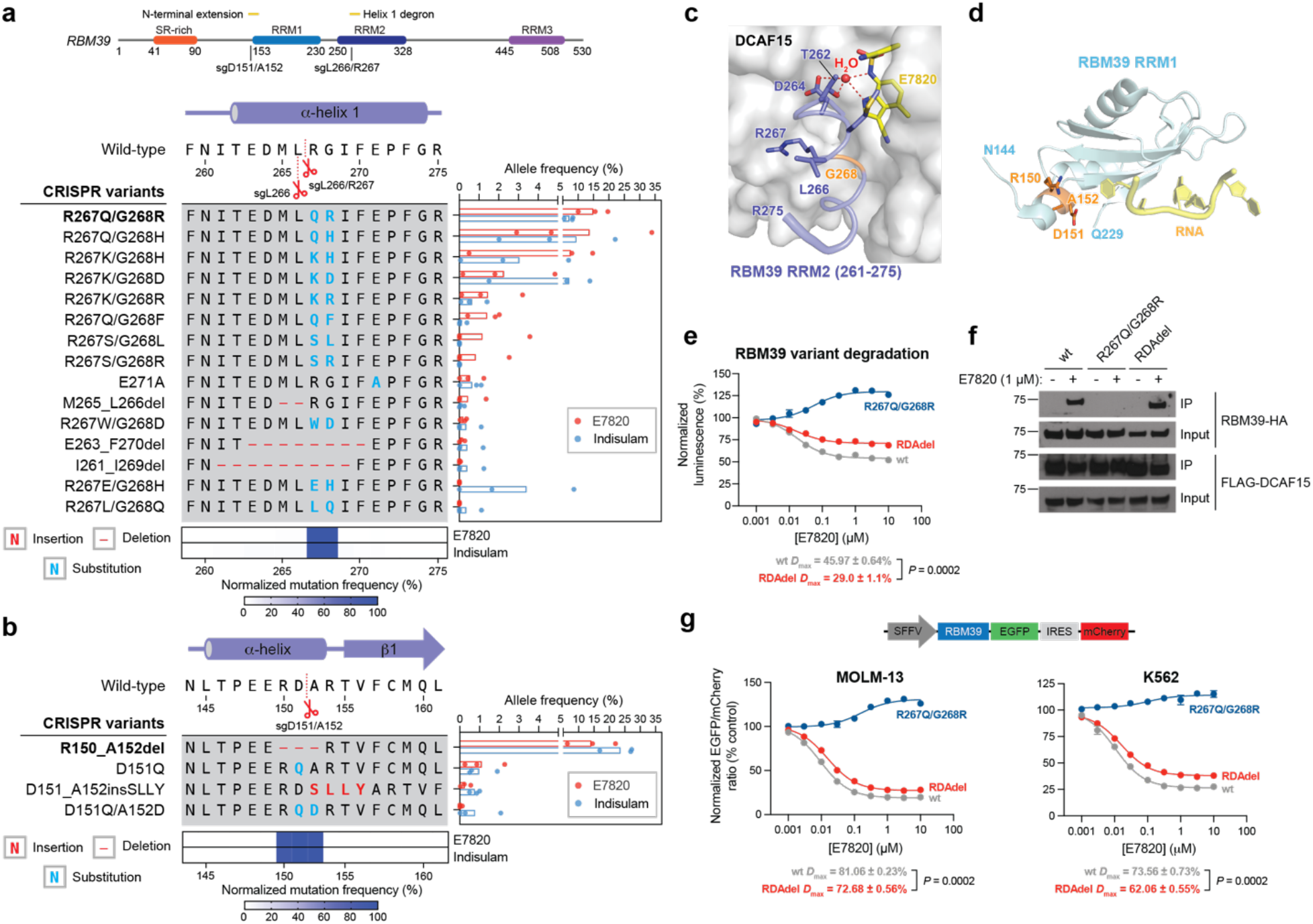
E7820 resistance mutations in different domains of RBM39 operate via distinct mechanisms. a) Left: Schematic shows the coding variants of the most abundant in-frame mutations enriched in the RRM2 helix 1 structural degron of RBM39 (>1% frequency in any condition). Right: Bar plot showing frequency (%, *x* axis) of each variant. Bars represent the mean across three replicate treatments and dots show the individual replicate values. Bottom: Heat map showing normalized mutational frequency (*y* axis, %) by sequence position (*x* axis). Mutational frequency was normalized as a percentage of the total frequency of the displayed variants. b) Left: Schematic shows the coding variants of the most abundant in-frame mutations enriched in the RRM1 N-terminal extension of RBM39 (>1% frequency in any condition). Right: Bar plot showing frequency (%, *x* axis) of each variant. Bars represent the mean across three replicate treatments and dots show the individual replicate values. Bottom: Heat map showing normalized mutational frequency (*y* axis, %) by sequence position (*x* axis). Mutation frequency was normalized as a percentage of the total frequency of the displayed variants. c) Structural view of the E7820-DCAF15-RBM39(RRM2) ternary complex, with key residues of RBM39 highlighted in blue. RBM39 G268 and a water molecule are highlighted in orange and red, respectively. Carbon atoms of E7820 are depicted in yellow. (PDB: 6UE5) d) Structural view of the RBM39 RRM1 domain (light blue), with key residues corresponding to the RDA deletion highlighted in orange (PDB:4YUD). RNA molecule from a CUGBP1 structure (PDB: 3NMR) is shown in yellow, overlaid and visualized by structural alignment. e) Dose-response curves for wt and mutant HiBiT-RBM39-HA cellular protein levels, as indicated by vehicle-normalized luminescence (*y* axis, %), in HEK293T cells treated with E7820 for 24 h. Data represent mean ± s.e.m. across three technical replicates. The *P* value was calculated using a two-sided Student’s *t*-test. One of two independent experiments is shown. f) Immunoblots showing co-IP of RBM39-HA wt and mutant variants with FLAG-DCAF15 after vehicle or E7820 treatment (1 µM, 4 h) in transiently transfected HEK293T cells. All cells were first pre-treated with MLN-4924 (1 µM, 2 h) prior to vehicle or E7820 treatment. Co-IP was performed using an anti-FLAG antibody. One of two independent replicates is shown. g) Top: Schematic of the fluorescent EGFP-IRES-mCherry degradation reporter vector. Bottom: Dose-response curves for wt and mutant RBM39 cellular protein levels, as indicated by vehicle-normalized EGFP to mCherry ratio (*y* axis, %), in MOLM-13 (left) or K562 (right) cells treated with E7820 for 24 h. Data represent mean ± s.e.m. across three technical replicates. The *D*_max_ ± s.e.m. and *P* values (two-sided Student’s *t*-test) are shown below. One of two independent experiments is shown for MOLM-13 cells, while one independent experiment was conducted for K562 cells.

This analysis revealed strong enrichment of diverse in-frame indel mutations within the GSPT1 structural degron in CC-885- and ZXH-1-161-treated conditions (**Fig 2a**). These mutations were predominantly centered around the sgK573 cut site, consistent with sgK573 being the most highly enriched sgRNA across both compound treatments, and frequently altered up to 7 amino acids spanning D571 to K577 with distinct amino acid deletions and insertions (**Fig 2a**, bottom panel). Consideration of the CC-885-CRBN-GSPT1 structure suggested that many of these mutations may impede ternary complex formation by modifying the conformation of the β-hairpin structural degron and consequently disrupting contacts between GSPT1, CC-885, and CRBN (**Fig 2b**).^10^ For instance, GSPT1 S574del removes a key serine residue that, together with D571, forms an ST-turn that stabilizes the β-hairpin (**Fig S2a**). In addition, several mutants, such as GSPT1 S574_K577del, altogether remove the critical G575 residue – the only residue within the β-hairpin degron conserved across all reported CRBN-IMiD neosubstrates.^10,13^ By contrast, mutations surrounding C568 are predicted to indirectly perturb the position of the β-hairpin by altering the upstream N-terminal sequences (C568_L569delinsQ) or by disrupting the ASX-motif involving D571 (C568_D571del) (**Fig S2b-c**). Lastly, mutations surrounding Q611 could not be detected, consistent with the lower resistance score of sgQ611 in comparison to sgC568 and sgK573. Corroborating these predictions, GSPT1 mutants in the presence of CC-885 failed to (1) degrade as assessed by a HiBiT lytic bioluminescence assay^34^ (**Fig 2c**) and (2) co-immunoprecipitate (IP) with CRBN in comparison to wild-type (wt) GSPT1 (**Fig 2d**). The diversity of resistance mutations that alter the sequence or position of the β-hairpin structural degron (vide infra) suggests that this region of GSPT1 can tolerate substantial sequence variation without compromising protein function essential for cell survival, and hence can serve as a hotspot for resistance mutations.

Likewise, for RBM39 we observed significant enrichment of in-frame mutations in the α-helical structural degron in RRM2 helix 1 across both E7820- and indisulam-treated conditions (**Fig 3a**). In contrast to GSPT1, the most prevalent mutations were double missense mutations primarily affecting R267 and G268 (**Fig 3a**, bottom panel), a previously identified hotspot for resistance.^14,16^ The tight packing of G268 against DCAF15 has been demonstrated in structural studies to preclude larger residues at this position without abrogating ternary complex formation (**Fig 3c**).^17,19^ Supporting this notion, RBM39 R267Q/G268R failed to degrade in the presence of E7820 in comparison to wt RBM39 (**Fig 3e**). Whereas the diversity of complex indels observed in the GSPT1 degron region suggests substantial mutational tolerance, the preponderance of double amino acid substitutions arising from extensive point mutations suggest stricter structural and/or functional constraints on the RBM39 degron. Altogether, these data spanning GSPT1 and RBM39 characterize the landscape of resistance mutations – comprising point substitutions and complex indels – that directly alter the structural degron while maintaining essential protein function.

### E7820 resistance mutations in different domains of RBM39 operate via distinct mechanisms

While the enrichment of sgRNAs targeting the structural degrons corroborate past findings, the enrichment of particular sgRNAs targeting regions outside of the RBM39 RRM2 helix 1 was unanticipated, as the binding affinity of RBM39 to DCAF15-sulfonamide predominantly depends on the RRM2 domain.^15,17,19^ Notably, sgD151/A152 is the second most highly enriched sgRNA in the *RBM39* CRISPR-suppressor scan for both sulfonamide treatments, suggesting that the resultant mutation(s) represents a major, competitive resistance mechanism on par with perturbing the structural degron (**Fig 1d**). Sequencing the amplicon surrounding sgD151/A152 revealed strong enrichment of RBM39 R150_A152del (abbreviated from here on as RDAdel) in the degrader-versus vehicle-treated pools (**Fig 3b**). The RDAdel mutation truncates an N-terminal α-helical extension that lies just outside of the annotated RRM1 domain (**Fig 3d**).

Due to its striking enrichment, we conducted a deeper investigation of RBM39 RDAdel. In contrast to the RRM2 helix 1 R267Q/G268R mutant, RBM39 RDAdel led to modest differences in RBM39 degradation versus wt RBM39 in a HiBiT assay conducted in HEK293T cells (**Fig 3e**). These differences in RBM39 degradation were predominantly characterized by a decreased level of maximal degradation (*D*_max_) at higher E7820 doses versus a change in half-maximal degradation concentration (DC_50_) (wt RBM39: DC_50_ = 17 nM, *D*_max_ = 46%; RBM39 RDAdel DC_50_ = 17 nM, *D*_max_ = 29%) (**Fig 3e, Fig S3a**). Notably, the differential *D*_max_ between wt and RDAdel RBM39 was dependent on expression levels of DCAF15, as overexpression of DCAF15 caused *D*_max_ to converge between the variants (**Fig S3b**). Co-IP experiments demonstrated that RBM39 RDAdel formed a ternary complex with DCAF15 and E7820 at comparable propensity as wt RBM39, while RBM39 R267Q/G268R completely failed to do so (**Fig 3f**). These results demonstrate that the RDA deletion does not fully abrogate E7820-DCAF15-RBM39 ternary complex formation and subsequent RBM39 degradation, suggesting that it may operate through a more intricate mechanism.

To corroborate the partial effects on *D*_max_, we evaluated RBM39 degradation using a fluorescent reporter system, in which wt or mutant RBM39 is fused in-frame with EGFP followed by an internal ribosome entry site (IRES) and mCherry (**Fig 3g**)._^13,27^_ Using this reporter, levels of RBM39 are directly correlated with EGFP fluorescence, which can be normalized to mCherry fluorescence to account for differences in reporter integrations and transcript expression levels. After lentiviral transduction of the reporter into MOLM-13 cells, levels of EGFP and mCherry fluorescence were assessed by flow cytometry after treatment with vehicle or E7820. As anticipated, cells expressing wt RBM39-EGFP exhibited a dose-dependent decrease in EGFP to mCherry ratio upon treatment with E7820 (DC_50_ = 9 nM, *D*_max_ = 81%) (**Fig 3g, Fig S4a**). By contrast, cells expressing RBM39-EGFP R267Q/G268R exhibited no decrease in EGFP to mCherry ratio even at the highest E7820 dose, as expected due to this mutant’s inability to form the DCAF15-RBM39 ternary complex (**Fig 3f**). However, cells expressing RBM39-EGFP RDAdel recapitulated a partial but significant rescue in degradation in comparison to wt RBM39-EGFP (RBM39-EGFP RDAdel: DC_50_ = 15 nM, *D*_max_ = 73%). Analogous effects were also observed in K562 and HEK293T cells (**Fig 3g, Fig S4b**), confirming our findings in multiple cell lines and across different degradation assays.

As ectopic expression of RBM39 RDAdel revealed modest differences in *D*_max_, we sought to characterize the RDA deletion in an endogenous context by generating clonal cell lines. We lentivirally transduced MOLM-13 cells with SpCas9 and sgD151/A152 and treated them with E7820 (1 μM) for 4 weeks, after which surviving cells were sorted, expanded, and genotyped. We identified clonal cell lines harboring homozygous RDAdel alleles, which we refer to as MOLM-13^*RDAdel*^ (**Fig S5a**). We confirmed that MOLM-13^*RDAdel*^ cells were resistant to treatment with E7820 and express RBM39 at levels comparable to wt MOLM-13 cells (**Fig 4a, Fig S5b**). Furthermore, immunoblotting after 24 hours of E7820 treatment revealed elevated levels of RBM39 in MOLM-13^*RDAdel*^ versus wt MOLM-13 cells at higher doses of E7820 tested (**Fig 4b**), consistent with the decreased *D*_max_ observed in ectopic expression experiments.

**Figure 4.**
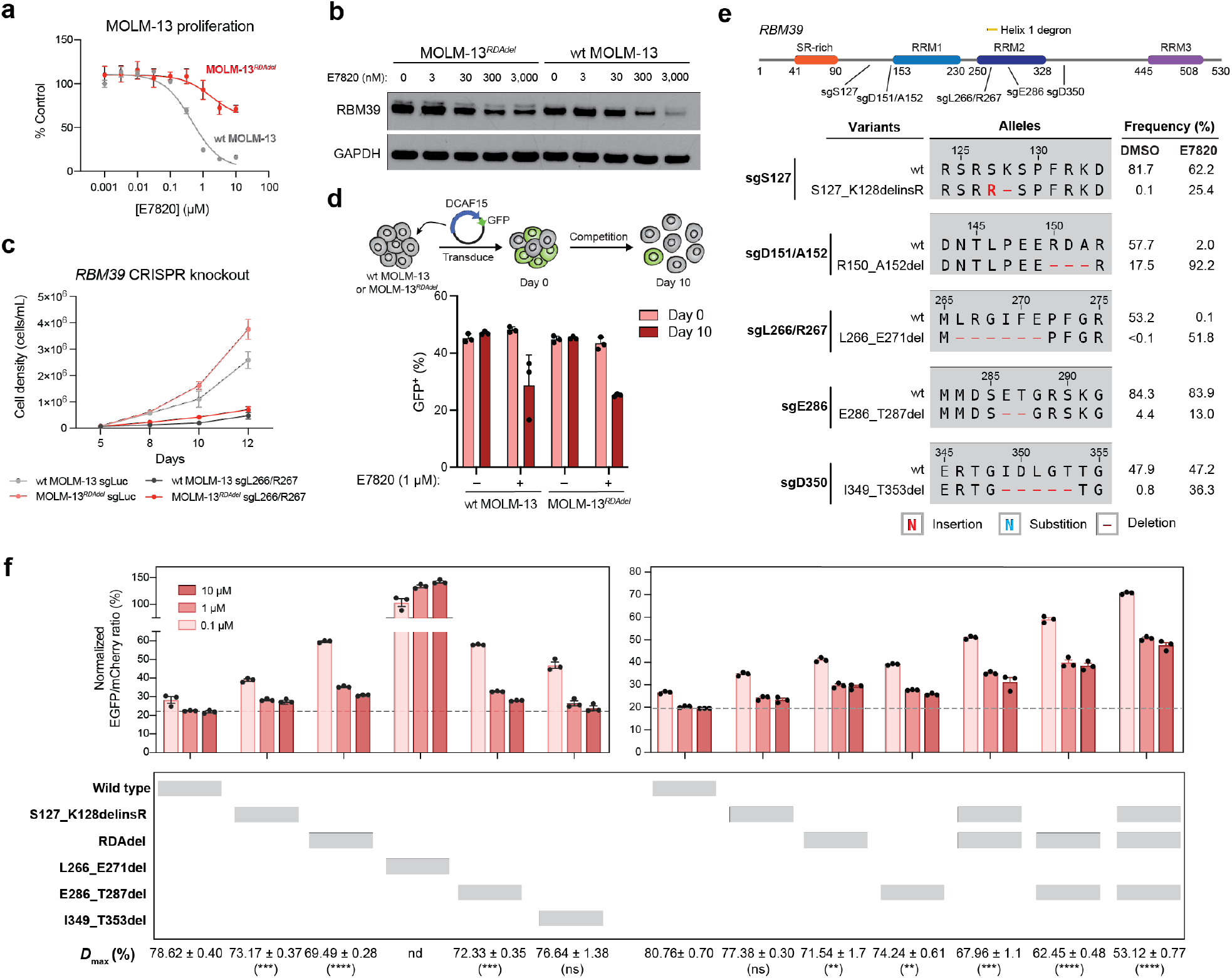
Mutations distal to the RBM39 RRM2 helix 1 structural degron alter maximum levels of RBM39 degradation to abrogate E7820 cytotoxicity. a) Dose-response curves for wt MOLM-13 and MOLM-13^RDAdel^ cell proliferation relative to vehicle-treated cells (*y* axis, % control) after E7820 treatment for 72 h. Data represent mean ± s.e.m. across three technical replicates. One of two independent experiments is shown. b) Immunoblots showing levels of RBM39 and GAPDH after vehicle or E7820 treatment for 24 h. One of two independent replicates is shown. c) Line graphs showing cell proliferation (*y* axis) over a time course (*x* axis) following lentiviral transduction of SpCas9 and sgRNAs targeting *luciferase* (sgLuc) or *RBM39* (sgL266/R267) into wt MOLM-13 and MOLM-13^RDAdel^ cells. Data represent mean ± s.e.m. across three technical replicates. One of two independent experiments is shown. d) Bar graphs showing fraction of GFP-positive cells (*y* axis) in a competition growth assay with non-transduced cells at day 0 and day 10 after treatment with either vehicle or 1 µM E7820 following lentiviral transduction of plasmid overexpressing DCAF15 and GFP in wt MOLM-13 and MOLM-13^RDAdel^. One of three independent replicates is shown. e) Schematic showing the coding variants of the most abundant in-frame RBM39 mutations enriched in E7820 treatment (1 µM) by each sgRNA tested. Variant frequencies in vehicle- and E7820-treatment conditions are indicated. f) Bar plots showing wt and mutant RBM39 cellular protein levels, as indicated by vehicle-normalized EGFP to mCherry ratio (*y* axis, %), in MOLM-13 cells treated with E7820 for 24 h. Data represent mean ± s.e.m. across three technical replicates. Dotted gray line indicates the mean signal of wt MOLM-13 treated with 10 µM E7820. Values for *D*_max_ ± s.e.m. are shown (right) with significance levels from a two-sided Student’s *t*-test comparing to wt RBM39 *D*_max_ indicated in parentheses (*P* < 10^−3^: ***; *P* < 10^−4^: ****; ns: not significant; nd: not determined). One of two independent experiments is shown. Full dose-response curves are shown in **Supplementary Figure S5d**.

Based on these results, we considered whether the RBM39 RDA deletion confers resistance by preventing depletion of RBM39 below a threshold level necessary to induce significant growth inhibition. Lowering RBM39 RDAdel levels by genetic depletion with CRISPR-Cas9 in MOLM-13^*RDAdel*^ led to growth inhibition, supporting the idea that substantial depletion of RBM39 RDAdel remains anti-proliferative in the MOLM-13^*RDAdel*^ cell line (**Fig 4c**). As expression of DCAF15 is correlated to *D*_max_ (**Fig S3b**), we considered whether ectopic overexpression of DCAF15 might re-sensitize MOLM-13^*RDAdel*^ to E7820. Indeed, both wt MOLM-13 and MOLM-13^*RDAdel*^ cells overexpressing DCAF15 exhibited growth inhibition upon E7820 treatment (**Fig 4d**). Taken together, our data support the possibility that even modest differences in target degradation can confer robust resistance to degraders and that mutations outside the ternary complex interface may be sufficient to achieve this.

### Several mutations distal to the RBM39(RRM2) structural degron decrease *D*_max_

Aside from sgD151/A152, we next considered whether other enriched sgRNAs targeting regions outside the RBM39 RRM2 helix 1 structural degron may generate resistance mutations that behave like the RDA deletion by partially decreasing *D*_max_ and impeding maximal RBM39 degradation. As we were unable to detect mutations at these distal sites by directly sequencing the pooled cells derived from the CRISPR-suppressor scan, we individually transduced selected enriched sgRNAs (RBM39 sgS127, sgE286, sgE343, sgD350) along with SpCas9 into MOLM-13 cells to identify their corresponding mutations. sgL266/R267 and sgD151/A152 were also transduced individually as comparators. Transduced cells were subsequently split and treated with E7820 (1 µM) or vehicle for 4 weeks and surviving cells were genotyped by targeted amplicon sequencing around the corresponding sgRNA cut sites.

With transduction of sgD151/A152 and sgL266/R267, we observed significant enrichment of in-frame mutations and concomitant depletion of the wt allele in the presence of E7820 versus vehicle control (**Fig 4e**). For sgL266/R267, in-frame variants constituted <0.5% of detected alleles under vehicle treatment, whereas the wt allele was highly prevalent at >50%. Under E7820 treatment, however, in-frame mutations and the wt allele represented >50% and <0.5% of detected alleles, respectively. These results suggest that RRM2 helix 1 variants may confer a significant fitness advantage to E7820 but may otherwise be rare and/or potentially deleterious in its absence (vide infra). By contrast, RDAdel was the predominant variant in cells transduced with sgD151/A152, comprising >90% and 17.5% of alleles in E7820- and vehicle-treatment, respectively. The prevalence of RDAdel in the control condition likely reflects its high predicted frequency as an editing outcome and limited effects on protein fitness and cell viability (**Fig S5c**).

In comparison to sgD151/A152, E7820-treatment led to more modest enrichment of in-frame alleles generated by sgS127, sgE286, and sgD350, consistent with these sgRNAs having lower resistance scores in the CRISPR-suppressor scan (**Fig 4e**). Mutations were not observed with sgE343 in this experiment. To assess possible effects on RBM39 degradation, we selected top enriched in-frame mutants generated by each sgRNA to evaluate with the RBM39-EGFP-IRES-mCherry reporter (**Fig 4f, Fig S5d**). As anticipated, complete rescue from E7820-induced degradation was observed with L266_E271del, which substantially alters the RRM2 helix 1 structural degron. By contrast, apart from I349_T353del, the remaining distal RBM39 mutants conferred partial resistance to E7820-induced degradation at levels similar to RDAdel (**Fig 4f, Fig S5d**). Notably, E286_T287del alters a β-hairpin within the RBM39 RRM2 domain formed by residues D284-R289 that may compromise a peripheral protein-protein interaction with DCAF15 (**Fig S5e**).^17–19^ Like RDAdel, S127_K128delinsR lies outside the RRM2 domain and is not structurally resolved.

We next considered if the distal mutations’ effects on *D*_max_ might be dependent or additive. RBM39 constructs containing two or three of these distal mutations exhibited significant cumulative decreases in *D*_max_ values of up to 25% (**Fig 4f, S5d**), showing that these mutations have additive effects and might operate independently of one another. Collectively, these findings show that several sites distal to the RBM39 structural degron, and in some cases distal to the known ternary complex interface altogether, can modulate *D*_max_ and the efficacy of target degradation. Furthermore, the observation that many distal site mutations can decrease *D*_max_ supports the notion that modest rescue of RBM39 levels is sufficient to confer resistance to E7820.

### Resistance mutation sites across TPD targets exhibit varying levels of sequence conservation and mutational constraint

Evolutionary conservation of protein sequences is a strong indicator of function. Consequently, protein sequence conservation can influence the accessible landscape of drug resistance-conferring mutations, as highly conserved sites (e.g., enzyme active sites) are typically more constrained by their functional importance and hence more difficult to mutate than less conserved sites. As a result, small molecules that bind or mechanistically involve less conserved sites may be more susceptible to the emergence of resistance mutations. Unlike orthosteric inhibitors, which typically modulate target activity and exploit the conserved structural features of active sites, molecular glue degraders are not necessarily dependent on neosubstrate function for efficacy. Thus, the mechanism of TPD may co-opt regions of the neosubstrate that are otherwise non-functional and hence may exhibit varying levels of mutational constraint. These considerations raise questions as to what factors shape the accessibility of neosubstrate resistance mutations.

Taking advantage of our CRISPR-suppressor scanning data spanning *GSPT1* and *RBM39*, we considered how sequence conservation and mutational constraint may influence the emergence and diversity of resistance mutations. To do so, we first calculated the conservation score of each residue in GSPT1 and RBM39 using ConSurf (**Fig 5a**), which estimates the relative conservation of each amino acid position (see Methods). As sequence conservation can vary substantially between adjacent residues and Cas9 generally mutates multiple amino acids around the cut site, we applied a LOESS regression to estimate per-residue conservation scores with respect to neighboring residues in the local region. As anticipated, this analysis highlighted the greater relative conservation of the well-defined protein domains versus the unstructured N-terminus and inter-domain linkers of each respective protein, where more negative ConSurf scores indicate higher levels of evolutionary conservation. In support of these calculations, our CRISPR-suppressor scanning data under vehicle-treatment showed preferential depletion of sgRNAs targeting more conserved regions (**Fig S6a-b**). The depletion of sgRNAs targeting functional protein regions has been previously demonstrated to indicate their essentiality, and consequently we refer to a sgRNA’s depletion in the vehicle condition as the “fitness score,”^35–37^ with lower scores corresponding to higher levels of essentiality.

**Figure 5.**
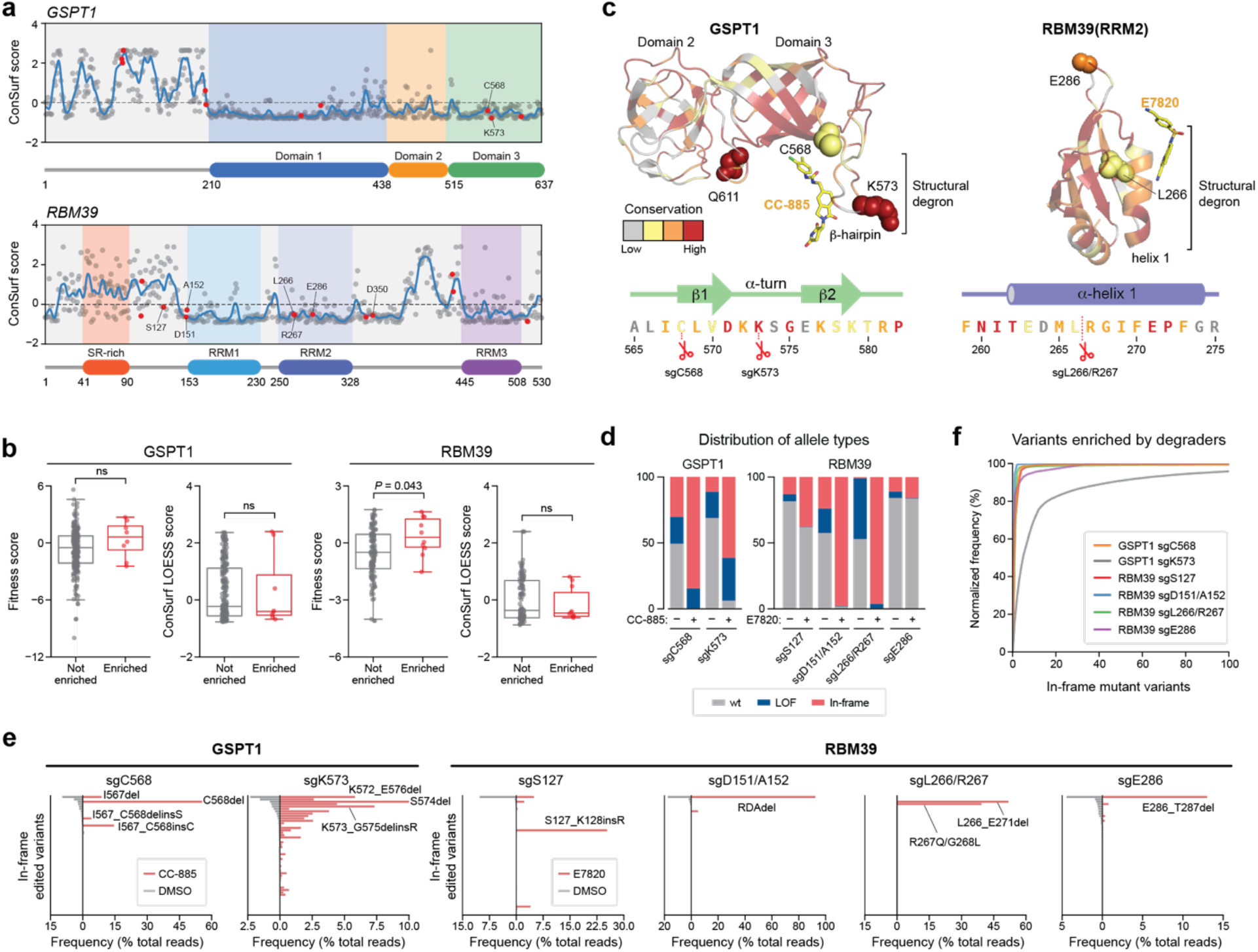
Resistance mutation sites across TPD targets exhibit low levels of sequence conservation. a) ConSurf conservation scores (*y* axis) of amino acid residues in *GSPT1* (top panel) and *RBM39* (bottom panel) shown as dots with the LOESS regression line in blue. Amino acids corresponding to enriched sgRNA cut site positions from the CRISPR-suppressor scanning are highlighted in red and key residues are labelled. b) Box plots with jitter showing fitness scores and ConSurf LOESS scores for non-enriched (gray, *n* = 230 for *GSPT1* and 119 for *RBM39*) or enriched (red, *n* = 9 for *GSPT1* and 10 for *RBM39*) sgRNAs. Fitness scores were calculated as the log_2_(fold-change sgRNA enrichment at week 4 under vehicle treatment versus the plasmid library) normalized to the mean of the negative control sgRNAs. sgRNAs were assigned ConSurf LOESS scores based on the amino acid corresponding to their predicted cut site positions; sgRNAs cutting between amino acids were assigned the mean of the flanking amino acids’ scores. Dots represent the fitness scores or corresponding amino acid ConSurf LOESS scores for individual sgRNAs. Two-sided *P* values were calculated with the Mann-Whitney test (ns: not significant). The box shows the median, 25^th^, and 75^th^ percentiles with whiskers denoting 1.5 × the interquartile range. c) Structural view of GSPT1(I440–P634) (left) and RBM39(RRM2) (right), with residues colored according to ConSurf conservation scores. The top three most conserved bins of ConSurf scores are colored in red, orange, and yellow, respectively, and the bottom six bins are colored in gray. sgRNAs enriched in the CRISPR-suppressor scan are depicted as spheres. Sequences corresponding to the approximate region around the structural degrons are shown below and colored according to ConSurf scores. d) Stacked bar plot showing the frequency distribution of variant types (*y* axis, % of total reads) after transduction of the indicated sgRNAs targeting *GSPT1* and *RBM39* and treatment with vehicle or drug molecules (see Methods) for 4 weeks. e) Bar plots showing variant frequencies (*x* axis, % of total reads) for the top 50 variants (*y* axis) generated by the indicated sgRNAs after treatment with vehicle (gray bars, left) or drug molecules (red bars, right) for 4 weeks. Variants are rank ordered on the *y* axis by decreasing frequency in vehicle treatment for each sgRNA. f) Cumulative plot showing the normalized variant frequency (*y* axis) for the 100 most abundant in-frame edited variants (*x* axis) for each indicated sgRNA after drug treatment for 4 weeks. Variants are rank ordered on the *x* axis by decreasing normalized frequency for each respective sgRNA condition. Variant frequency was normalized to the total frequency of all in-frame edited variants.

We next assessed the sgRNA fitness scores and conservation of positions in GSPT1 and RBM39 implicated in mediating resistance, focusing on enriched sgRNAs that had resistance scores ≥ 2 s.d. above the mean of the negative controls in either degrader condition. In RBM39 and GSPT1, top enriched sgRNAs by resistance score had similar fitness scores to non-enriched sgRNAs and were close to 0, the mean of the negative controls, indicating that they are functionally neutral and not likely targeting highly essential positions (**Fig 5b**). By ConSurf, these positions exhibited comparable or slightly greater conservation than those targeted by non-enriched sgRNAs. Altogether, these data suggest that resistance mutations to degraders can occur at sites that are not highly conserved relative to other residues.

Beyond evolutionary conservation, we next sought to directly assess the permissible mutational landscape across resistance sites validated in our study, including those affecting the (1) GSPT1 β-hairpin structural degron, (2) RBM39 RRM2 helix 1 structural degron, and (3) RBM39 distal positions. Although the neosubstrate structural degrons are both relatively conserved (**Fig 5c**), degrader treatments more strongly jackpot sgRNAs targeting the GSPT1 β-hairpin (sgC568, sgK573) than sgRNAs targeting the RBM39 RRM2 helix 1 (sgL266, sgL266/R267) within their respective CRISPR-suppressor scans, as the RBM39 distal mutations can presumably compete effectively with the RRM2 helix 1 mutations despite their partial rescue phenotype (**Fig 4e-f**). We therefore reasoned that the GSPT1 β-hairpin and RBM39 distal positions may tolerate more mutational variation than the highly structured RBM39 RRM2 helix 1, permitting more diversity of mutations in these regions that do not abrogate essential function. To explore this notion further, we first individually transduced GSPT1 sgC568 and sgK573 – sgRNAs that target the β-hairpin – along with SpCas9 into MOLM-13 cells, treated with either vehicle or CC-885 for 4 weeks, and then genotyped the surviving cellular pools by targeted amplicon sequencing. We then compared these allele frequency data for GSPT1 with those acquired previously for RBM39 sgL266/R267, sgS127, sgD151/A152, and sgE286 – sgRNAs that target the RRM2 helix 1 and distal positions, respectively (**Fig 4e**).

We first considered alleles identified under the vehicle conditions. While frequencies of in-frame edited alleles typically ranged from 10% to 30% across both *RBM39* and *GSPT1*, the total percentage of in-frame edited alleles generated by RBM39 sgL266/R267 was significantly lower (<1%) (**Fig 5d**), suggesting that the RRM2 helix 1 cannot tolerate mutational variation. It is unlikely that this difference in in-frame edited alleles is due to major discrepancies in sgRNA cutting efficiencies, as the fraction of total edited alleles for sgL266/R267 is second highest amongst the sgRNAs evaluated (**Fig 5d, Fig S5c**). We next scrutinized the distribution of the in-frame variants under vehicle conditions (**Fig 5e, Fig S5c**). sgRNAs targeting the GSPT1 β-hairpin – in particular sgK573 – led to a wider spread of in-frame variant distributions than sgRNAs targeting *RBM39*. Altogether, these results suggest that resistance sites to degraders exhibit a wide range of mutational constraint under normal growth conditions, and that the GSPT1 β-hairpin can tolerate mutational variation to a higher extent than positions in RBM39 despite its sequence conservation.

We next evaluated allele frequencies identified in the degrader-treated conditions. Across all sgRNAs evaluated, in-frame mutant alleles were enriched by degrader treatment (**Fig 5d**), and as anticipated, this enrichment was greatest for top-scoring sgRNAs in the CRISPR-suppressor scans (i.e., GSPT1 sgC568, GSPT1 sgK573, RBM39 sgD151/A152, RBM39 sgL266/R267). We considered how the distribution of in-frame variants may change between vehicle- and degrader-treated conditions. Under degrader treatment, the distributions of in-frame mutations generally become more skewed and rarer variants can be highly selected for (**Fig 5e**,**f, Fig S5c**), consistent with not all mutations robustly conferring resistance. However, the level of skewing is highly variable. In particular, in-frame variant distributions generated by introduction of RBM39 sgL266/R267 and sgD151/A152 are dominated by 1-2 mutants each under E7820-treatment. For RBM39 sgL266/R267, this jackpotting supports the idea that RBM39 RRM2 helix 1 mutations are highly selected for and constrained. By contrast, the jackpotting observed with RBM39 sgD151/A152 likely reflects the prevalence of RDAdel as a favorable editing outcome that is both well-tolerated and selected (**Fig 4e**). Other RBM39 distal position mutations were also highly selected for by E7820, albeit to a lesser extent than those generated by sgL266/R267 and sgD151/A152 (**Fig 5e**,**f, Fig S5c**).

Strikingly, mutagenesis of the GSPT1 β-hairpin by sgK573 and, to a lesser extent, gC568 led to wider spread distributions of in-frame variants in comparison to mutagenesis of RBM39 (**Fig 5e**,**f**). Altogether, these data suggest that the GSPT1 β-hairpin can accommodate many mutations, a large fraction of which can robustly confer resistance (**Fig S5c**). Moreover, many of these in-frame mutations identified in the GSPT1 β-hairpin involve complex indel mutations altering variable stretches of multiple amino acids (**Fig 2a**) in contrast to the predominance of point mutations or smaller deletions observed in RBM39 upon E7820-treatment (**Fig 3a, Fig 4e**).

As a result, despite its high sequence conservation, the GSPT1 degron can tolerate substantial mutational variation, in contrast to the highly conserved and mutationally constrained degron of RBM39. This mutational constraint imposed on the RBM39 degron, in tandem with the modest rescue effect required to restore growth, likely enabled the emergence of diverse resistance mutations across distal positions of RBM39 (i.e., S127_K128delinsR, RDAdel, E286_T287del). Altogether, our analysis highlights how various factors can constrain or enable the accessibility of resistance mutations to degraders and cooperate to ultimately shape neosubstrate-specific mutational landscapes.

## Discussion

Whereas resistance mutations to occupancy-driven inhibitors are well-studied and fall broadly into several archetypal classes (e.g., drug binding disrupting, enzyme activating), the analogous mutational landscape for molecular glue degraders remains poorly defined owing to their unique mode of action.^2,4^ To address these challenges, here we systematically profiled the landscape of resistance mutations afforded by CRISPR-mutagenesis across two distinct TPD neosubstrates, GSPT1 and RBM39. We demonstrate that CRISPR-suppressor scanning can rapidly identify mutations that confer resistance to molecular glue degraders, and that most of these mutations disrupt the structural degron. Such mutations are consistent with structural data and reinforce the notion that high-grade resistance mutations may disrupt ternary complex formation by perturbing either small molecule-protein interactions, protein-protein interactions, or both. Based off these observations, we expect that CRISPR-suppressor scanning will be a convenient approach to study potential neosubstrate surfaces and their interactions involved in ternary complex formation, especially in the absence of structural information or where multiple binding modes may possibly occur (e.g., heterobifunctional degraders).

While mutations at the molecular glue interface can readily disrupt ternary complex formation and TPD, we also identified unexpected resistance mutations in RBM39 that are distal to the structural degron. Through closer investigation, we show that some distal mutations decrease the depth of maximal protein degradation (i.e., *D*_max_), thereby preventing sufficient target depletion that is necessary for growth inhibition to occur. Moreover, these impacts on *D*_max_ can be additive, showing how multiple, independent distal structural alterations may be sufficient to significantly alter target degradation. Interestingly, increased DCAF15 expression diminishes the impact of RDAdel on *D*_max_, suggesting that mutations that modestly decrease *D*_max_ might have stronger effects in cells with lower E3 ligase expression. Of note, the threshold of target degradation necessary for phenotypic response may be highly context-dependent, which may further influence the potential of these “depth-altering” resistance pathways. For example, studies investigating BCL6-degraders suggest that very high levels of BCL6 degradation are necessary to achieve tumor regression^39^ and, in these instances, mutations that only modestly influence *D*_max_ may be sufficient to confer resistance. These observations highlight the advantage of investigating resistance mutations within their endogenous protein contexts, as dosage effects may be prevalent. Altogether, our findings suggest that distal mutations can be sufficient to decrease *D*_max_ and degrader efficacy, raising the possibility that distal post-translational modifications, alternative isoforms, or even binding partners may have similar effects as well.

Lastly, by integrating CRISPR-suppressor scanning data spanning *GSPT1* and *RBM39*, we demonstrate that mutational tolerance of the structural degron is a primary driver of the overall landscape of degrader resistance mutations within neosubstrates. Neosubstrates like GSPT1, where the β-hairpin structural degron is not mutationally constrained, may permit the formation of mutational “hotspots,” with an array of diverse mutations concentrated within a small region of the protein. On the other hand, neosubstrates like RBM39, where the α-helical structural degron is highly constrained, can lead to the emergence of degrader resistance mutations across numerous sites both within and distal to the structural degron, despite their weaker effects on *D*_max_. While these mutations are largely context-specific and generated by CRISPR-Cas9, which favors formation of indels, these results highlight the utility in profiling mutational constraint directly, which may complement evolutionary conservation analysis. More broadly, our analysis highlights how the interplay of relative mutational constraint across putative sites of resistance can shape divergent outcomes in the overall landscape of degrader resistance mutations.

In conclusion, systematic identification of drug resistance-conferring alleles through approaches like CRISPR-suppressor scanning can illuminate neosubstrate requirements that are necessary for chemically-induced dimerization and for TPD to drive effective phenotypic responses. Notably, many secondary neosubstrate features beyond the structural degron are not well understood and typically involve flexible yet potentially functional regions that are not structurally resolved, highlighting the utility of this approach. Deeper investigation of the mutants identified in these types of studies might uncover additional resistance mechanisms. We anticipate that the strategy developed here will also be broadly applicable for the study of TPD across different types of degraders (e.g., PROTACs, autophagy-targeting chimeras), neosubstrates, and E3 ligase systems, and will be informative for the design and optimization of degraders.

## Materials and Methods

### Cell culture and lentiviral production

MOLM-13 cells were a gift from M.D. Shair (Harvard University). HEK293T cells were a gift from B.E. Bernstein (Massachusetts General Hospital). 293FT and K562 cells were obtained from Thermo Fisher Scientific and ATCC, respectively. All cell lines were cultured in a humidified 5% CO_2_ incubator at 37 °C and routinely tested for mycoplasma (Sigma-Aldrich). All media were supplemented with 100 U ml^−1^ penicillin and 100 µg ml^−1^ streptomycin (Gibco) and FBS (Peak Serum). MOLM-13 and K562 cells were cultured in RPMI-1640 (Life Technologies) supplemented with 10% FBS. HEK293T and 293FT cells were cultured in DMEM (Gibco) supplemented with 10% FBS, 6 mM L-glutamine (Gibco), 0.1 mM MEM non-essential amino acids (Gibco), and 1 mM sodium pyruvate. For lentivirus production, the transfer plasmids of interest were co-transfected with *GAG/POL* and *VSVG* plasmids into 293FT cells using FuGENE HD (Promega) according to manufacturer’s protocol. Media was exchanged after 6-8 h and the viral supernatant was collected 48 h after transfection and sterile-filtered (0.45 μm). MOLM-13 and K562 cells were transduced by spinfection at 1,800 × *g* for 1.5 h at 37 °C with 5 μg ml^-1^ and 1 μg ml^-1^ polybrene (Santa Cruz Biotechnology), respectively. 48 h post-transduction, cells were selected with 1 μg ml^-1^ and 2 μg ml^-1^ puromycin (Thermo Fisher Scientific) respectively for 10 days.

### Chemical reagents

Compounds were stored at –80 °C in 100% DMSO. The vehicle condition represents 0.1% (v/v) DMSO treatment. Indisulam and E7820 were purchased from Sigma-Aldrich (≥98% purity by HPLC) and Cayman Chemical (≥98% purity by HPLC), respectively. MLN-4924 was purchased from MedChemExpress (≥98% purity by NMR). CC-885 was purchased from MedKoo Biosciences (98% purity by HPLC) and ZXH-1-161 was a gift from N. S. Gray (Stanford University).

### Cell growth assays

MOLM-13 cells were seeded at a cell density of 3 × 10^5^ cells/mL in a 96-well plate in triplicate with drug or vehicle treatments. Cell viability was assessed by flow cytometry after 3 days of drug or vehicle treatment using Helix NP NIR viability dye (BioLegend). Dose-response curves were fitted through interpolation using GraphPad Prism v.7 nonlinear regression fit ([inhibitor] versus normalized response–variable slope).

### Pooled sgRNA library cloning and CRISPR-suppressor scanning experiments

The *RBM39* and *GSPT1* tiling libraries contained all sgRNAs targeting the coding sequence (*NP_909122*.*1* and *NP_002085*.*3*) with MIT specificity scores above 25 (CRISPOR) and were ordered as an oligo pool from Twist Biosciences. These sgRNA sequences are listed in the **Supplementary Table 1**-**2**. The oligo pool was cloned into pLentiCRISPR.v2 as previously described.^40^ pLentiCRISPR.v2 was a gift from F. Zhang (Addgene 52961, Broad Institute). Lentiviral particles carrying the resulting sgRNA library were generated as described above and titered according to the published procedure.^40^ Cells (13-15 × 10^7^) were transduced at a multiplicity of infection < 0.3 and subsequently selected with 1 μg ml^-1^ puromycin for 10 days. The cells were then split into pools and treated with the drug(s) or vehicle in triplicate. The cells were passaged every 3-4 days and seeded at a density of 0.3-0.4 × 10^6^ cells/mL. The cells transduced with the *RBM39* sgRNA library were treated with 1 μM of sulfonamide analogs (indisulam or E7820) while the cells transduced with the *GSPT1* sgRNA library were treated with increasing concentrations of drugs (CC-885 and ZXH-1-161) according to the growth curve to produce a consistent impact on cell proliferation. Briefly, for CC-885, the cells were initially treated at 1 nM for a week, with doses escalating each subsequent week to 2.5 nM, 5 nM, and 10 nM over 4 weeks. For ZXH-1-161, the cells were initially treated at 85 nM for a week, with doses escalating each subsequent week to 500 nM, 750 nM, and 1000 nM over 4 weeks. Genomic DNA was isolated from the drug-and vehicle-treated cells at the specified time points using the QIAamp DNA Blood Mini Kit (Qiagen). The sgRNA composition of the population from each replicate was PCR amplified and barcoded followed by next-generation sequencing on a MiSeq (Illumina) using 150-cycle, single-end reads as previously described.^41^

### CRISPR-suppressor scanning data analysis

All data processing and analysis were performed using Python v.3.8.3 (www.python.org). Raw sequencing data were processed as previously described.^32,42^ In brief, reads were counted by identifying the 20-nt sequence downstream of the ‘CGAAACACCG’ prefix and mapped against a reference file containing all library sgRNA sequences with 0 mismatch allowance. sgRNAs with 0 reads in the plasmid library were excluded from the analysis. Read counts were then converted to reads per million, log_2_-transformed after adding a pseudocount of 1 to each sgRNA, and then normalized by subtracting the log_2_-transformed sgRNA counts in the plasmid library. sgRNA enrichment scores were calculated by averaging across replicates for each condition and normalized by subtracting the mean enrichment scores of the negative control sgRNAs. This value as calculated in the vehicle-treated conditions is referred to as the “fitness score.” sgRNA “resistance scores” were calculated using sgRNA enrichment scores in the drug-treated conditions. sgRNAs were classified as “enriched” if their resistance scores were greater than the mean + 2 standard deviations of the negative control sgRNAs. sgRNAs were mapped to their respective protein amino acid positions using the genomic coordinates of the predicted cut site in the *RBM39* (*NP_909122*.*1*) and *GSPT1* (*NP_002085*.*3*) coding sequences. sgRNAs were assigned to a single amino acid if the cut site fell within a codon or assigned to the two flanking amino acids if the cut site fell between codons.

### Individual sgRNA validation experiments and genotyping data analysis

Raw sequencing data were processed and aligned to *RBM39* and *GSPT1* using CRISPResso2 (v.2.0.40) to identify genomic variants and quantify allele frequencies.^43^ An in-house Python pipeline was used to classify and characterize variants at the protein level. Variants were classified as ‘in-frame’ if the indel size was a multiple of three and did not span an intron-exon junction. In-frame variants were then globally re-aligned to the reference coding sequence at the nucleotide level with a custom codon-aware implementation of the Needleman-Wunsch algorithm using the ‘PairwiseAligner’ module of Biopython (v.1.7.8), trimmed, and translated into their corresponding protein variants.^44^ Variants were classified as ‘loss-of-function’ if the indel size was not a multiple of three (i.e., frameshift), led to a premature stop codon (i.e., nonsense), or disrupted canonical splice site positions (the 2 nt immediately flanking each exon). Editing outcome predictions for individual sgRNAs were obtained using the inDelphi web server (https://indelphi.giffordlab.mit.edu) in single mode with K562 as the cell-type.^45^

### Plasmids

Open reading frames (ORFs) for RBM39 (wt and variants), GSPT1 (wt and variants), CRBN, and DCAF15 were cloned into pcDNA3.1 vectors using Gibson cloning (New England Biolabs). HA and FLAG tags were introduced through the primers used for Gibson cloning. RBM39 and GSPT1 proteins with N-terminal HiBiT tags were cloned into an expression vector with the herpes simplex virus thymidine kinase (TK) promoter using Gibson cloning. RBM39 wt and variants were cloned into the Artichoke reporter plasmid (a gift from B. Ebert, Addgene 73320) using Golden Gate cloning.

### Lytic HiBiT detection assay

HEK293T cells (0.8 × 10^6^) were plated in a 6-well plate. After 24 h, cells were transfected with 50 ng of plasmids encoding the HiBiT-tagged proteins using FuGENE HD (Promega). Cells were trypsinized (Gibco) after 24 h and 20,000 cells were plated per well in triplicate in white, opaque 96-well plates (Corning). Cells were allowed to attach for 24 h and then treated with various concentrations of compounds or 0.1% vehicle for 6 h (CC-885) or 24 h (E7820). An equal volume of Nano-Glo HiBiT reagent (Promega) containing the lytic buffer, substrate and the LgBiT protein was then added according to the manufacturer’s protocol. The plate was incubated at room temperature for 10 min with shaking at 350 rpm before measuring end point luminescence on a SpectraMax i3x microplate reader (Molecular Devices).

### Fluorescent degradation reporter assay

Lentiviral particles carrying the respective constructs in the Artichoke EGFP-IRES-mCherry reporter vector were produced and used to transduce MOLM-13 and K562 cells as described above. 48 h after transduction, cells were selected with appropriate puromycin concentration for 3-5 days. The selected cells were then treated with various concentrations of E7820 or 0.1% vehicle for 24 h. EGFP and mCherry fluorescence were measured on a NovoCyte 3000RYB flow cytometer (Agilent) after drug or vehicle treatment. The geometric mean of the ratio of EGFP to mCherry fluorescence was calculated for each sample using the NovoExpress software (v. 1.5.0, Agilent). The ratios for the individual drug-treated samples were normalized to the ratios of the vehicle-treated samples.

### Immunoblotting

MOLM-13 cells were plated in triplicate at 30,000 cells per well in a 96-well plate. After treatment with various concentrations of E7820 or 0.1% vehicle for 24 h, cells were harvested and washed with PBS (Corning) once. Cells were then lysed in RIPA buffer (Boston BioProducts) supplemented with 1× Halt Protease Inhibitor (Thermo Fisher Scientific). Total protein concentration in the clarified lysates was determined using the BCA Protein Assay Kit (Thermo Fisher Scientific). Samples were prepared for SDS-PAGE followed by immunoblot analysis according to standard procedures. For immunoblotting, anti-Caper (Bethyl laboratories, A300-291A, 1:15,000), anti-HA (Cell Signaling Technology, 3724S, 1:1,000), anti-FLAG (Cell Signaling Technology, 2368S, 1:1000), anti-CRBN (Novus Biologicals, NBP1-91810, 1:500), anti-CRBN (Cell Signaling Technology, 71810, 1:1,000), anti-GAPDH (Santa Cruz Biotechnology, sc-47724), and anti-β-actin antibody (Sigma-Aldrich, A1978, 1:5,000) were used.

### Co-immunoprecipitation

HEK293T (3 × 10^5^) cells were plated into a 6-well plate and transfected 24 h post-plating with 400 ng pcDNA3.1-HA-RBM39 and 800 ng pcDNA3.1-DCAF15 expression vectors using FuGENE HD (Promega). 48 h post-transfection, cells were pre-treated with 1 µM MLN-4924 for 2 h. After 2 h, cells were treated with either vehicle or 1 µM E7820 for 4 h. Cells were then harvested and washed with PBS. Cells were lysed in lysis buffer (50 mM NaCl, 50 mM NaH_2_PO_4_, 50 mM sodium citrate, 20 mM HEPES pH 7.4, 1% NP-40, 5% glycerol) supplemented with vehicle or E7820 (1 µM) and the protein concentration of the cell lysate was quantified as described above. Cell lysates containing 500 µg of protein were then incubated with Protein G Dynabeads (Thermo Fisher Scientific) and 2 µg anti-FLAG M2 antibody (Sigma-Aldrich) overnight with rotation at 4 °C followed by washing with the lysis buffer three times. Samples were then prepared for SDS-PAGE and analyzed by immunoblotting as described above. For co-immunoprecipitation of CRBN with GSPT1 variants, HEK293T (3 × 10^6^) cells were seeded in a 10 cm plate a day prior to transfection. The cells were transiently transfected with plasmids expressing GSPT1-HA wild type and variants (6 µg) using lipofectamine 3000 according to manufacturer’s protocol (Thermo Fisher Scientific). After 48 h, the cells were treated with 1 µM MLN-4924 for 3 h followed by treatment with either vehicle or 10 µM CC-885 for 2h. Cells were then harvested, washed with PBS, snap frozen and stored at –80 °C until further use. Cells were then lysed with 500 µL lysis buffer (50 mM Tris pH 7.5, 50 mM NaCl, 0.5% NP-40, 10% glycerol, 1X Halt protease inhibitor cocktail (Thermo Fisher Scientific), 1X EDTA and 1X PhosphoSTOP (Roche) supplemented with 1 µM MLN-4924 and vehicle or 10 µM CC-885 and the protein concentration was quantified as described above. Cell lysates containing 1 mg of protein were incubated with anti-HA antibody (1:100) for 1 h at 4 °C followed by incubation with Protein G Dynabeads (Thermo Fisher Scientific) for 1 h at 4 °C. The beads were then washed with lysis buffer supplemented with with 1 µM MLN-4924 and vehicle or 10 µM CC-885, three times. Samples were then prepared for SDS-PAGE and analyzed by immunoblotting as described above.

### Sequence conservation analysis

Sequence conservation scores were obtained for RBM39 and GSPT1 using the ConSurf web server (https://consurf.tau.ac.il).^46^ ConSurf scores were computed using the ConSeq method.^47^ In brief, ConSurf homologues were collected from the UniRef90 database using HMMER and up to 250 homologues with 70% to 99% sequence identity were sampled from the list of unique hits for multiple sequence alignment, phylogenetic tree construction, and conservation scoring. ConSurf scores are normalized such that the mean score across all input residues is zero and the standard deviation is one. ConSurf scores are relative, with the lowest scores representing the most conserved positions in the input sequence. As sequence conservation can exhibit significant variation between adjacent residues, LOESS regression was performed on the ConSurf scores to assess the overall conservation profile of each residue with respect to neighboring residues. LOESS regression was performed using the ‘lowess’ function of the statsmodels package (v.0.12.1) in Python with frac = (10 AA/L), it = 0.

### Statistical methods

All statistical tests described were performed as two-sided tests. Pearson coefficients and significance values were calculated using the ‘stats.pearsonr’ function of the SciPy package (v.1.6.0) in Python. Other statistical parameters including the exact value and definition of *n*, the definition of center, dispersion, precision measures (e.g., mean ± s.d. or s.e.m.), and statistical significance are reported in figures and figure legends. Data for protein degradation assays and cell proliferation assays were graphed and fit to sigmoidal curves by non-linear regression (GraphPad Prism). For degradation assays, the DC_50_ (IC_50_), *D*_max_ (100 – bottom asymptote), and corresponding s.e.m. values were determined from the sigmoidal fit.

### Protein computational modelling

Modeling of select GSPT1 mutants was performed using Molecular Operating Environment (Chemical Computing Group). Using the structure of wt GSPT1 complexed to CC-885, DDB1, and CRBN (PDB: 5HXB), each 5-7 residue stretch containing the desired mutation was modeled using the Loop Modeler application (RMSD limit = 1, Loop limit = 100, Energy window = 10). Where possible, PDB loop searching was preferred over de novo loop searching in identifying candidate loops. For each mutant, the candidate obtained with the lowest MM/GBVI energy was selected as the final model.

## Data Availability

Data are provided in the main text and figures, supplementary figures (**Fig S1-S5**) and tables (**Supplementary Table**).

## Code Availability

Custom python scripts for data analysis of CRISPR-suppressor scanning and single sgRNA-mutagenesis experiments are available upon request.

## Author Contributions

P.M.G. and K.C.N. contributed equally. P.M.G. and B.B.L. conceived the project. P.M.G., K.C.N., and B.B.L designed the experiments. P.M.G. performed the CRISPR-suppressor screens and single guide experiments for RBM39 and GSPT1. K.C.N. performed computational analysis. P.M.G. and C.S. cloned and performed the co-immunoprecipitation studies. P.M.G. characterized single cell clones, performed cell growth assays, and degradation assays using western blot. P.M.G. cloned HiBiT-tagged constructs and performed the HiBiT degradation assay. M.Y. and P.M.G. cloned and performed the protein degradation Artichoke reporter assay in K562 and MOLM-13. K.C.N. performed the protein degradation Artichoke reporter assay in HEK293T. J.M.L. synthesized select compounds used in the study. N.Z.L. performed computational protein modelling. S.M.H performed initial literature review. B.B.L., P.M.G., and K.C.N. wrote the manuscript. B.B.L. supervised the study and held overall responsibility of the study.

## Acknowledgements

The authors acknowledge the Gray lab for providing ZXH-1-161. The authors acknowledge Ceejay Lee for assistance in designing sgRNA libraries and members of the Liau lab for comments on the manuscript.

---

**Supplementary Figure S1.**
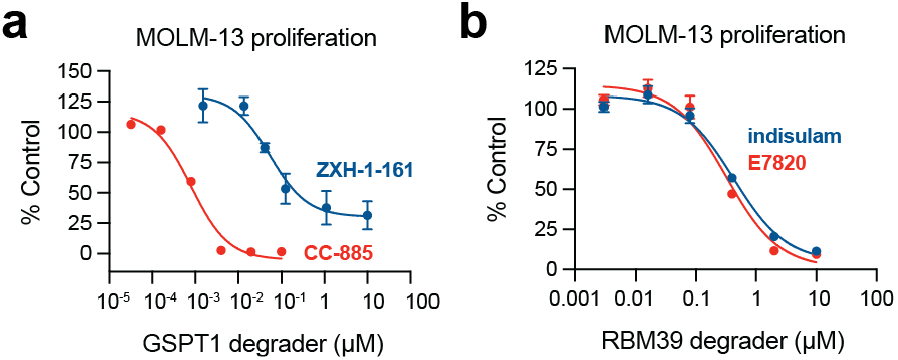
CRISPR-suppressor scanning of GSPT1 and RBM39. a) Dose-response curves for MOLM-13 cell proliferation relative to vehicle-treated cells (*y* axis, % control) after E7820 or indisulam treatment for 72 h. Data represent mean ± s.e.m. across three technical replicates. Experiment performed once. b) Dose-response curves for MOLM-13 cell proliferation relative to vehicle-treated cells (*y* axis, % control) after CC-885 treatment (72 h) or ZXH-1-161 treatment (48 h). Data represent mean ± s.e.m. across three technical replicates. Experiment performed once.

**Supplementary Figure S2.**
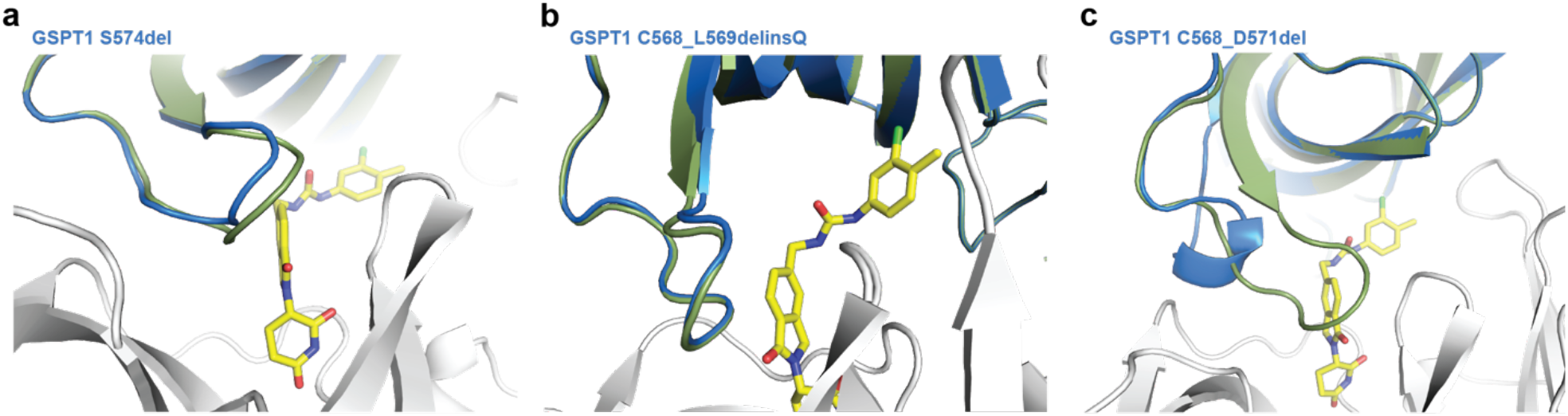
Protein computational modeling of selected CC-885 resistant GSPT1 mutants. Structural views of wt GSPT1 (green, PDB: 5HXB) overlaid with modeled GSPT1 resistant mutants (blue) in complex with CC-885 (yellow) and CRBN (white), showing changes in loop structure caused by the indicated indel mutations. GSPT1 S574del is shown in **a**, GSPT1 C568_L569delinsQ is shown in **b**, and GSPT1 C568_D571del is shown in **c**.

**Supplementary Figure S3.**
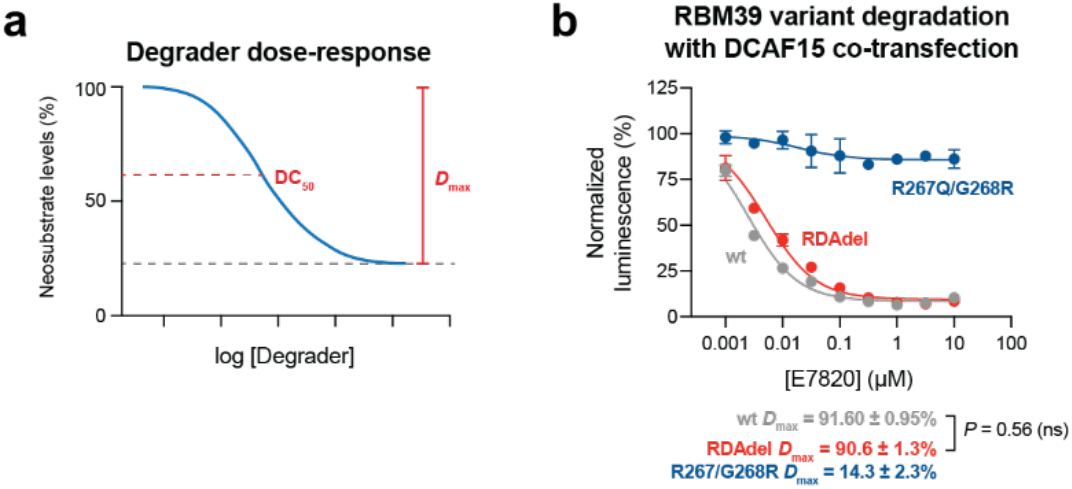
Degradation assay data for RBM39 variants. a) Schematic of a generic sigmoidal dose-response curve illustrating neosubstrate degradation (*y* axis, % protein) as a function of degrader concentration (*x* axis), with key parameters (DC_50_ and *D*_max_) denoted. b) Dose-response curves for wt and mutant HiBiT-RBM39-HA cellular protein levels, as indicated by vehicle-normalized luminescence (*y* axis, %), in HEK293T cells co-transfected with a DCAF15 expression plasmid (pcDNA3.1-DCAF15) and treated with E7820 for 24 h. Data represent mean ± s.e.m. across three technical replicates. The *D*_max_ ± s.e.m. and *P* values (two-sided Student’s *t*-test, ns: not significant) are shown below. One of two independent experiments is shown.

**Supplementary Figure S4.**
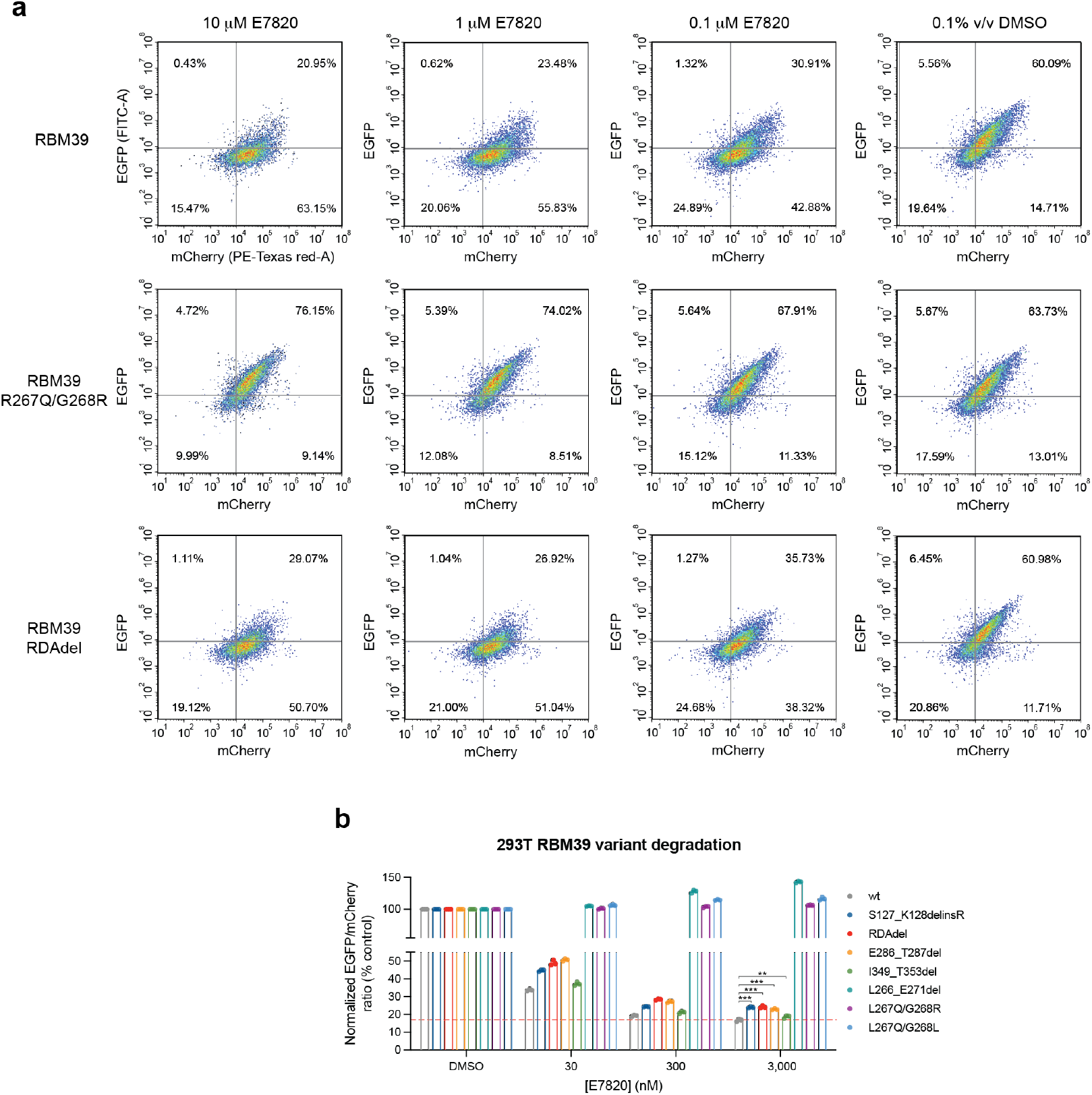
EGFP-IRES-mCherry fluorescent reporter assay for monitoring RBM39 variant degradation. a) Flow cytometry pseudocolor density plots showing EGFP (*y* axis) and mCherry (*x* axis) fluorescence for wt RBM39 (top), RBM39 R267Q/G268R (middle), and RBM39 RDAdel (bottom) after vehicle or E7820 treatment for 24 h. Populations were gated on mCherry-positive cells using the gating strategy in **Supplementary Figure S4c**. One of two independent experiments shown. b) Bar plots showing wt and mutant RBM39 cellular protein levels, as indicated by vehicle-normalized EGFP to mCherry ratio (*y* axis, %), after treatment with E7820 for 24 h in HEK293T cells stably expressing the RBM39-EGFP-IRES-mCherry fluorescent reporter. Data represent mean ± s.e.m. across three technical replicates. Dotted red line indicates the mean signal of wt MOLM-13 treated with 3 µM E7820. Significance levels comparing the degradation levels of the indicated RBM39 mutants to wt RBM39 at 10 μM E7820 are shown (*P* < 10^−2^: **; *P* < 10^−3^: ***, two-sided Student’s *t*-test). One of two independent experiments is shown.

**Supplementary Figure S5.**
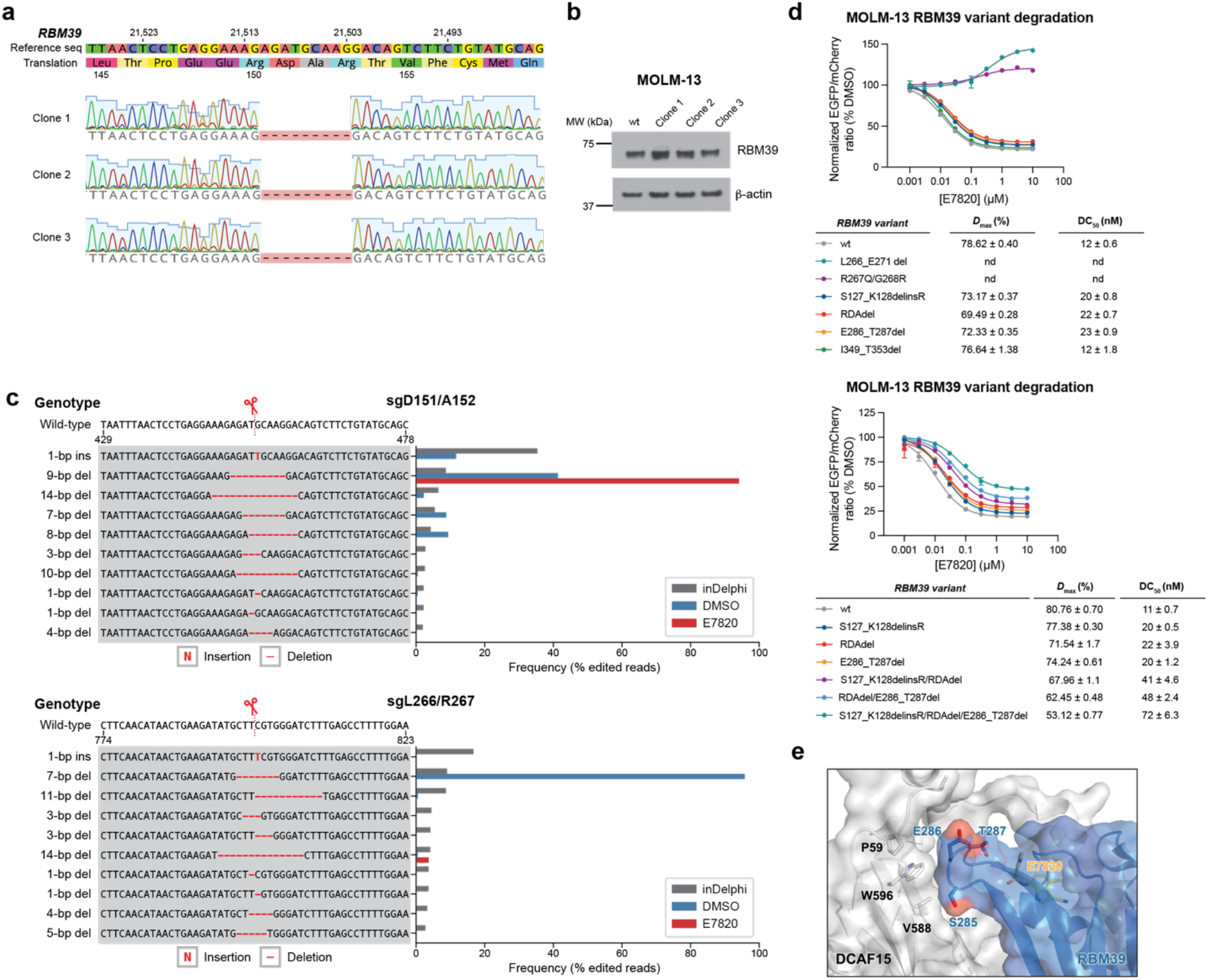
Characterization of MOLM-13^*RDAdel*^ and RBM39 variants identified by single sgRNA CRISPR mutagenesis. a) Schematic depicting the coding mutations and genotypes identified in the MOLM-13^*RDAdel*^ clonal cell lines. b) Immunoblots showing that RBM39 are expressed at comparable levels in wt MOLM-13 and MOLM-13^*RDAdel*^ clonal cell lines. One of two independent experiments is shown. c) Left: Schematic showing the top 10 editing outcomes predicted by inDelphi for gD151/A152 (top) and gL266/R267 (bottom). Right: Bar plot showing frequency (%, *x* axis) of each editing outcome (*y* axis) as predicted by inDelphi (gray) or as observed in MOLM-13 cells after vehicle (blue) or E7820 (red) treatment for 4 weeks. d) Dose-response curves for wt and mutant RBM39 cellular protein levels, as indicated by vehicle-normalized EGFP to mCherry ratio (*y* axis, %), in MOLM-13 cells treated with E7820 for 24 h. Top and bottom panels represent different experiments. Data represent mean ± s.e.m. across three technical replicates. Values for *D*_max_ ± s.e.m. and DC_50_ ± s.e.m. are tabulated (nd: not determined). One of two independent experiments is shown. e) Structural view of the E7820-DCAF15-RBM39(RRM2) ternary complex depicting the interaction between the RBM39(RRM2) (blue) D284–R289 hairpin and DCAF15 (grey), with E7820 shown in yellow (PDB: 6UE5).

**Supplementary Figure S6.**
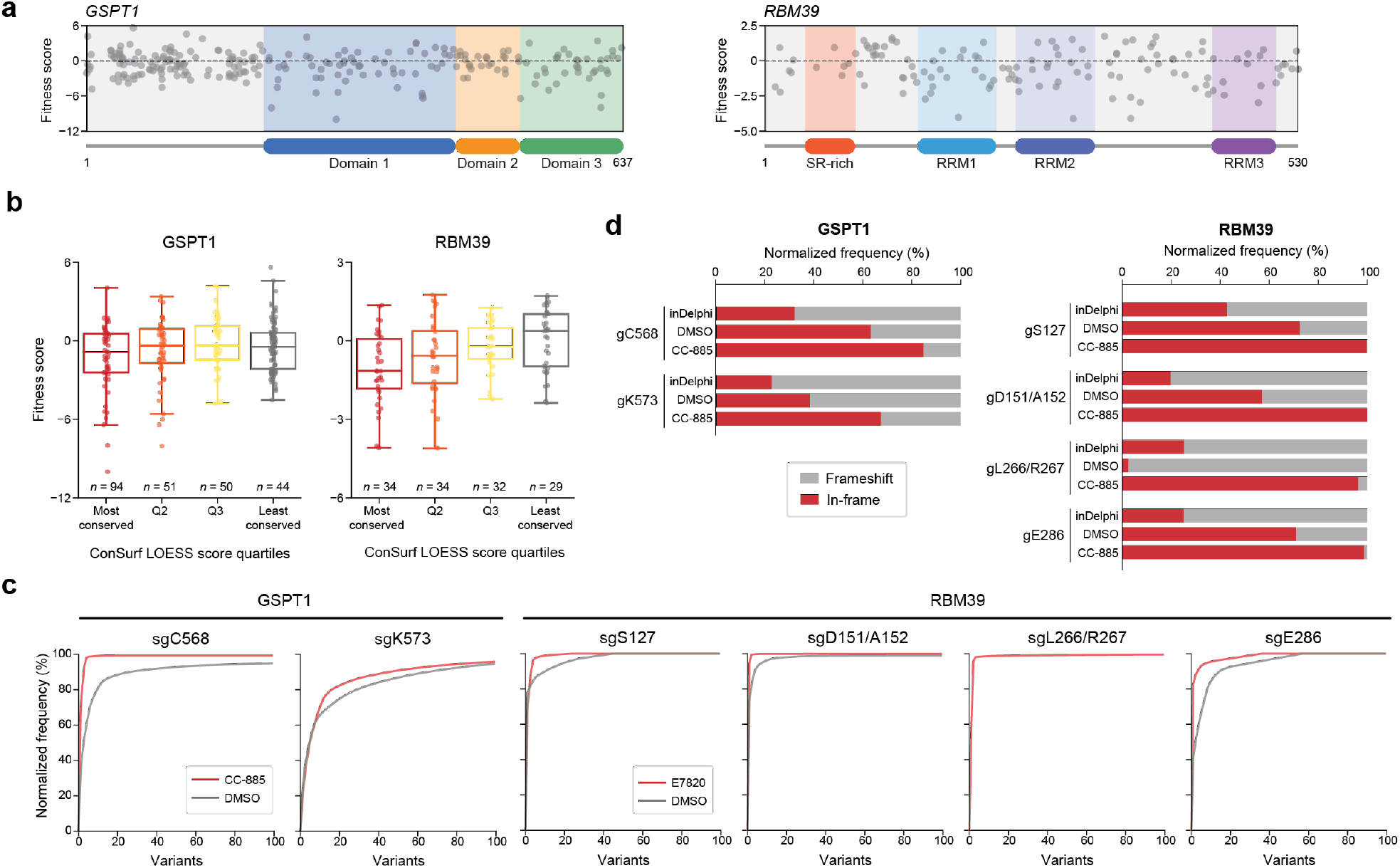
Analysis of GSPT1 and RBM39 fitness and CRISPR mutagenesis. a) Scatter plot showing sgRNA fitness scores (*y* axis) in MOLM-13 for CRISPR-scanning of *GSPT1* (left, *n* = 239) or *RBM39* (right, *n* = 129). Fitness scores were calculated as the log_2_(fold-change sgRNA enrichment at week 4 under vehicle treatment versus the plasmid library) normalized to the mean of the negative control sgRNAs (*n* = 22 for GSPT1 and 80 for RBM39). The sgRNAs are arrayed by amino acid position in the respective CDS on *x* axis corresponding to the position of the predicted cut site. When the sgRNA cut site falls between two amino acids, both amino acids are denoted. Data points represents mean value across three replicate treatments. Protein domains are demarcated by the colored panels. b) Box plots with jitter showing fitness scores for sgRNAs targeting *GSPT1* (left) and *RBM39* (right) grouped by the ConSurf LOESS score quartile. sgRNAs were assigned ConSurf LOESS scores based on the amino acid corresponding to their predicted cut site positions; sgRNAs cutting between amino acids were assigned the mean of the flanking amino acids’ scores. Dots represent individual sgRNAs. The box shows the median, 25^th^, and 75^th^ percentiles with whiskers denoting 1.5 × the interquartile range. c) Cumulative plots showing the normalized variant frequency (*y* axis, %) for the 100 most abundant in-frame edited variants (*x* axis) for each indicated sgRNA after vehicle or drug treatment (see Methods) for 4 weeks. Variants are rank ordered on the *x* axis by decreasing normalized frequency for each respective sgRNA condition. Variant frequency was normalized to the total frequency of all in-frame edited variants. The plot for RBM39 gL266/R267 under vehicle treatment is not shown since in-frame edited variants comprise <1% of the total frequency. d) Stacked bar plots showing the normalized frequency distribution of frameshift (gray) and in-frame (red) variant types (*y* axis, % of all editing outcomes) for the indicated sgRNAs as predicted by inDelphi or as observed in MOLM-13 after vehicle or drug treatment for 4 weeks. Variant type frequency was normalized to the total frequency of all edited variants. Note that inDelphi does not predict editing outcomes containing substitutions or insertions greater than 1-nt.

